# A semi-variance approach to visualising phylogenetic autocorrelation

**DOI:** 10.1101/2021.05.21.445056

**Authors:** M. J. Noonan, W. F. Fagan, C. H. Fleming

## Abstract

1. Comparing traits across species has been a hallmark of biological research for centuries. While inter-specific comparisons can be highly informative, phylogenetic inertia can bias estimates if not properly accounted for in comparative analyses. In response, researchers typically treat phylogenetic inertia as a form of autocorrelation that can be detected, modelled, and corrected for. Despite the range of methods available for quantifying the strength of phylogenetic autocorrelation, no tools exist for visualising these autocorrelation structures.
2. Here we derive variogram methods suitable for phylogenic data, and show how they can be used to straightforwardly visualise phylogenetic autocorrelation. We then demonstrate their utility for three empirical examples: sexual size dimorphism (SSD) in the Musteloidea, maximum per capita rate of population growth, *r*, in the Carnivora, and brain size in the Artiodactyla.
3. When modelling musteloid SSD, the empirical variogram showed a tendency for the variance in SSD to stabilise over time, a characteristic feature of Ornstein-Uhlenbeck (OU) evolution. In agreement with this visual assessment, model selection identified the OU model as the best fit to the data. In contrast, the infinitely diffusive Brownian Motion (BM) model did not capture the asymptotic behaviour of the variogram and was less supported than the OU model. Phylogenetic variograms proved equally useful in understanding why an OU model was selected when modelling *r* in the Carnivora, and why BM was the selected evolutionary model for brain size in the Artiodactyla.
4. Because the variograms of the various evolutionary processes each have different theoretical profiles, comparing fitted semi-variance functions against empirical semi-variograms can serve as a useful diagnostic tool, allowing researchers to understand why any given evolutionary model might be selected over another, which features are well captured by the model, and which are not. This allows for fitted models to be compared against the empirical variogram, facilitating model identification prior to subsequent analyses. We therefore recommend that any phylogenetic analysis begin with a non-parametric estimate of the autocorrelation structure of the data that can be visualized. The methods developed in this work are openly available in the new `R` package `ctpm`.

## Introduction

Comparing traits across species has been a hallmark of biological research for centuries (Harvey & Pagel, 1991; Garland *et al*., 2005). Indeed, inter-specific comparisons have yielded some of the most important biological advances, including the theory of evolution by natural selection (Darwin, 1859), allometric scaling rules (Bergman, 1848; Rensch, 1950; Jetz, 2004; Hirt *et al*., 2017; Noonan et al., 2020), the metabolic theory of ecology (Brown *et al*., 2004), and theories on the evolution of sociality (Lukas & Clutton-Brock, 2013; Noonan *et al*., 2015). While inter-specific comparisons can be highly informative, as early as Darwin (1859) it was recognised that the characteristics of newly evolved species are based on modifications of traits inherited from ancestors. This inheritance will limit the differences in traits between closely related taxa, especially if only a short amount of time has passed, a phenomenon known as ‘phylogenetic inertia’ (Blomberg & Garland Jr, 2002). In his seminal paper, Felsenstein (1985) showed how phylogenetic inertia can be viewed as as a form of statistical autocorrelation that could result in biased estimates and misleading significance if not properly accounted for in comparative analyses. Felsenstein (1985) effectively translated the concept of phylogenetic inertia into a statistical problem and provided a path forward for correcting for autocorrelation-induced biases in comparative analyses using statistical approaches.

The idea that phylogenetic inertia should leave a statistically identifiable autocorrelation structure in species trait data that could be modelled was transformative, and methods for modelling phylogenetic autocorrelation have evolved substantially over recent decades (e.g., Martins & Hansen, 1997; Abouheif, 1999; Butler & King, 2004; Harmon *et al*., 2008; Revell *et al*., 2008; Blomberg *et al*., 2020). Researchers now routinely model phylogenetic autocorrelation when making inter-specific comparisons (Abouheif & Fairbairn, 1997; Noonan *et al*., 2015; Johnson *et al*., 2017), and/or quantify the strength of phylogenetic autocorrelation to infer evolutionary processes (Morales, 2000; Kellermann *et al*., 2012; Herrera, 2020). These approaches primarily rely on modelling evolution according to Gaussian stochastic processes, typically some form of Brownian Motion (Einstein, 1905), Ornstein-Uhlenbeck processes (Uhlenbeck & Ornstein, 1930), or, more recently, non-Gaussian stochastic processes (Blomberg *et al*., 2020). Models are fit via maximum likelihood estimation (Harmon *et al*., 2008; Revell, 2012; Clavel *et al*., 2015), and the best model is chosen via standard model selection procedures (Burnham & Anderson, 2002). While existing tools for working with phylogenetic data have yielded novel insight into evolutionary processes (e.g., Furness *et al*., 2021; Smaers *et al*., 2021), a challenge of the conventional workflow is that it offers no way to visualise the autocorrelation structure of the data, nor to assess whether any of the candidate models look like the data. This is a notable limitation as the ability to visualise autocorrelation is crucial for understanding the underlying evolutionary process and determining what stochastic model the data best suggest. This stands in stark contrast to the numerous tools available for quantifying the strength of phylogenetic autocorrelation, such as Moran’s I (Moran, 1950), Pagel’s λ (Pagel, 1999), Abouheif’s C_mean_ (Abouheif, 1999), and Blomberg’s K (Blomberg *et al*., 2003) (reviewed in: Münkemüller *et al*., 2012). As a result of this limitation, researchers often rely on colouring the branch tips or lengths of phylogenetic trees based on trait values (e.g., Revell, 2013). While this can serve as a useful visual tool, it provides no information on the underlying stochastic process by which the trait may be evolving.

Here we show how semi-variograms offer both a novel approach for visualising phylogentic autocorrelation and a solution to the model diagnostic problem. Semi-variograms were originally developed to describe the degree of spatial dependence of random fields in geostatistics, as it was well known that geological samples taken taken close together in space would be more similar to one another than samples taken farther apart (Matheron, 1963). While originally developed to describe spatial autocorrelation, semi-variograms have also been extended to describe the temporal autocorrelation of stochastic processes (Fleming *et al*., 2014). Because the different stochastic processes used to model phylogenetic autocorrelation all have different theoretical variograms (Fleming *et al*., 2014), empirical semi-variograms can provide a useful diagnostic tool for checking a model’s fit (Pérez-Barbería *et al*., 2004). Here, we extend semi-variograms to the needs of phylogenetic autocorrelation, and develop confidence intervals on the estimated semi-variances. We then demonstrate the utility of the method for three empirical examples: i) the evolution of sexual size dimorphism in the carnivoran super-family Musteloidea (Noonan *et al*., 2016); ii) maximum per capita rate of population growth in the Carnivora (Fagan *et al*., 2013); and iii) brain size in the Artiodactyla (Haarmann, 1975; Oboussier, 1979).

## Materials and methods

### Semi-variograms for phylogenetic data

There are methods other than the variogram for visualising autocorrelation structures than can be applied to phylogenetic trait data (e.g., phylogenetic correlograms; Diniz-Filho, 2001), but we focus here on semi-variograms as they are the only known nonparametric autocorrelation estimator that can handle the highly irregular time intervals that typify phylogenetic tree data (see also Fleming *et al*., 2014).

Let *X_i_*(*t*) denote the trait *x* of the ancestor of species *i* at time *t* in the past, which all share the same stochastic process distribution, but evolve independently after bifurcation. Under an assumption of stationarity, the semi-variance function at lag *τ* can be estimated via any weighted average of the form

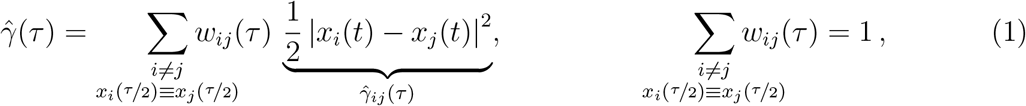

where the sum runs only over species that last shared a common ancestor at time *τ*/2 in the past, and where the second constraint fixes the expectation value 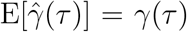. The above form of estimator is unbiased and asymptotically consistent, but its variance will be determined by our choice of weights *w_ij_*(*t*).

Optimal weights will minimize the variance

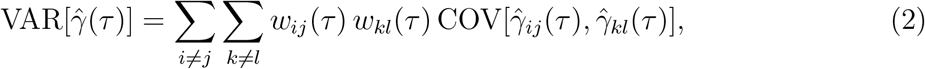

under constraint (1). This, however, requires an estimate of the model. First, at a given lag *τ*, let us represent the above quadratic form in matrix notation as

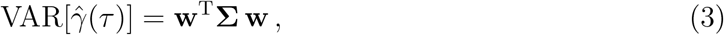

where the indices of the block-vector **w** are *ij* and the indices of the block-matrix **∑** are *ij*; *kl*. When including constraint (1), the Lagrangian is

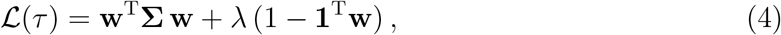

where **1** is the vector of all 1s. The optimal weights are, via straightforward calculation, given by

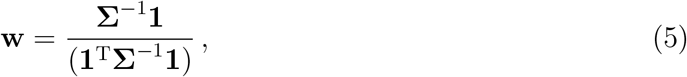

which requires inverting the covariance matrix **∑**. This, in turn, requires **∑** to be correctly positive definite (PD). Therefore, we cannot reliably optimize our weights based on a non-parametric estimate of the variogram, because it will not be ensured to provide a PD covariance function. Instead, we can rely on parametric assumptions, as all valid parametric models will always be PD. Moreover, we do not require all model parameters, as any overall constant (equivalent to the variance) will cancel out in (5). Given the stationary assumption, we only require the correlation matrix, **C**.

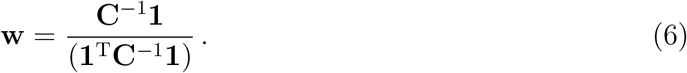

Here, our weights optimized to minimize variance can occasionally be negative, which can lead to slightly negative estimates when the true value is close to zero. This can be remedied by introducing the inequality constraints *w_ij_*(*τ*) ≥ 0, which turns (5) into a quadratic programming (QP) problem that can be solved to obtain a non-negative estimate of less optimal variance (Turlach, 2019). In that case, the above relations still hold if the resultant QP weights are non-negative.

### Independent and Identically Distributed (IID)

If the phylogenetic process is Independent and Identically Distributed (IID), then it is sufficient to consider the correlation matrix **C** ∝ **∑**, where the diagonal of **C** is 1 and the off-diagonal is 1/4 if species pair (*i, j*) and (*k, l*) share one species in common and 0 otherwise.

### Brownian Motion (BM)

If the phylogenetic process is BM, then it is sufficient to consider the correlation matrix **C** ∝ **∑**, where the diagonal of **C** is 1 and the off-diagonal is the squared proportion of time lag *τ* during which the backward-in-time-forward-in-time tip-branch-tip trajectories {*x_i_*(0) → *x_ij_*(*τ*/2) → *x_j_*(0)} and {*x_k_*(0) → *x_kl_*(*τ*/2) → *x_l_*(0)} correspond to the same species, where *x_i_*(*τ*/2) = *x_j_*(*τ*/2) = *x_ij_*(*τ*/2) (Fig. 1).

**Figure 1:**
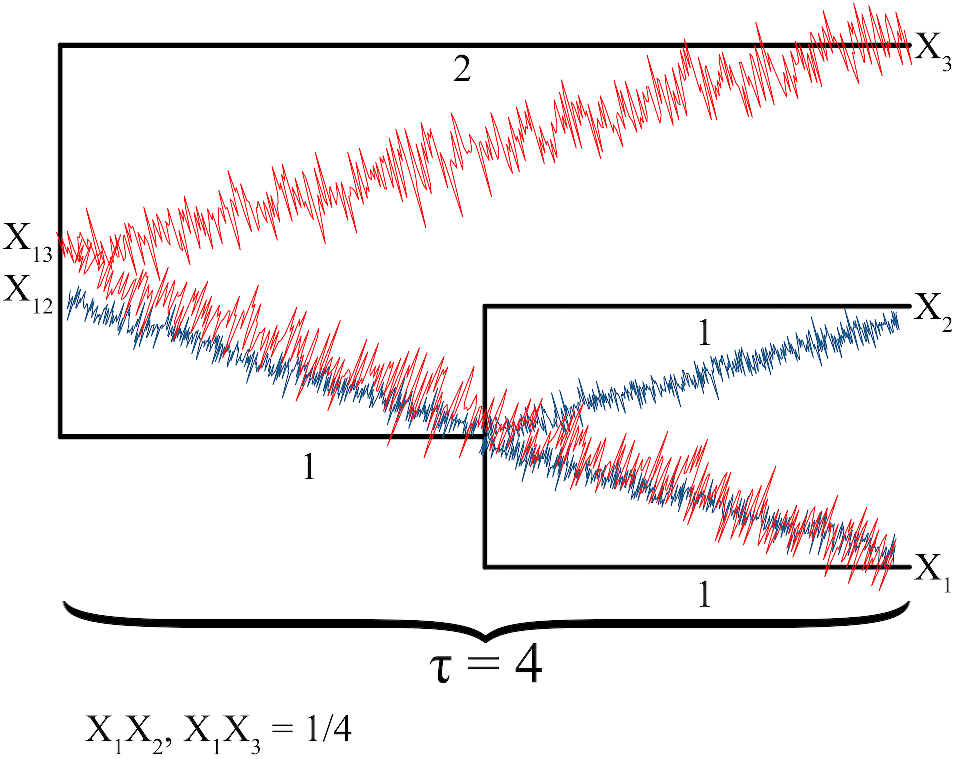
Schematic diagram depicting the calculation of the BM weights for the species pair (1, 2) and (1,3).

### Ornstein-Uhlenbeck (OU)

Optimal weights for an OU process can be derived, but there will be dependence on the unknown timescale parameter, which is why we do not consider such cases here.

### Confidence intervals

Using the same weights as before, and assuming the correlation structure to be correct, the variance of the variogram is given by

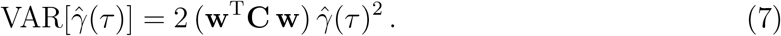

Variances are strictly positive and range between 0 and ∞, so as an improvement over normal confidence intervals, which can include negative values, we summarise the uncertainty in 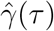 with χ^2^ statistics.

### Time-lag gridding

The time lags for phylogenetic variograms are calculated based on the topologies and branch lengths of the supplied phylogenetic trees, and the structure of a particular tree will dictate what pairwise time lags are possible. These trees represent snapshots of the evolutionary processes, and will almost always result in irregular time series. Although irregularity in the data is acceptable, coarsening the variogram according to time-lag bin widths can make variograms more straightforward to interpret. Here we identify the time-lag bins using Gaussian mixture model (McLachlan & Basford, 1988) and kmeans (MacQueen *et al*., 1967) clustering implemented in the `R` package `clusterR` (ver 1.2.4, Mouselimis, 2021), with the number of clusters, *n*, given as 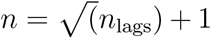, and initial estimates of the centroids given by the regular grid points running along the range of time-lags. This number of bins was selected as in the limit of a perfectly regularly time series, the number of bins would reduce back to the conventional grid.

The methods developed in this work are openly accessible in the new `R` package `ctpm`, available at https://github.com/NoonanM/ctpm.

### Semi-variance functions of evolutionary models

We next express IID, BM, and OU models in terms of their semi-variance functions. We selected these three processes as they are the models most frequently used in practice (e.g., Pérez-Barbería *et al*., 2004; Fagan *et al*., 2013; Furness *et al*., 2021; Smaers *et al*., 2021). We also provide a biological interpretation of the parameters of these three semi-variance functions. A diagram of how these semi-variance functions relate to different tree configurations and patterns of traits assumed under each model is shown in Figure 2.

**Figure 2:**
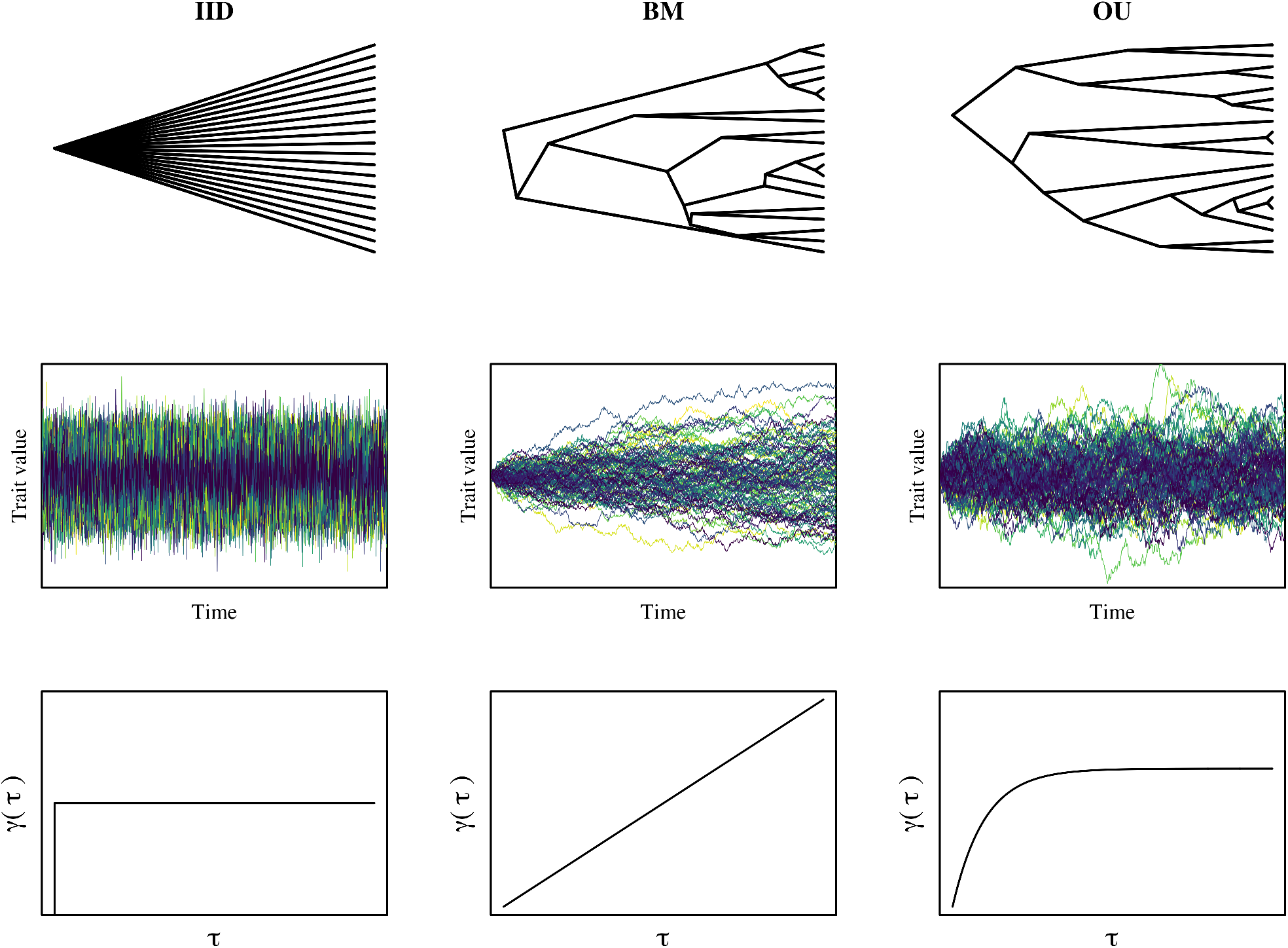
Examples of Independent and Identically Distributed (IID), Brownian Motion (BM), and Ornstein-Uhlenbeck (OU) phylogenetic models and their semi-variograms. The first row depicts the form of the phylogenetic tree assumed under each processes, the second the underlying stochastic processes, and the third the theoretical semi-variance functions. In particular, note how the BM and OU phylogenetic trees are difficult to interpret, whereas the semi-variance functions can be easily distinguished.

### IID: No phylogenetic inertia

Under an IID model of evolution, the values of traits observed in lineage *i* at one point in time *x_i_*(*t*) are independent of traits at any other time *x_i_*(*t*’), with *t* ≠ *t*’ and evolve according to a white noise process. In other words, there is no phylogenetic autocorrelation present under this evolutionary model and the semi-variance is therefore not dependent on the time-lag *τ*, but is simply given by the steady-state variance

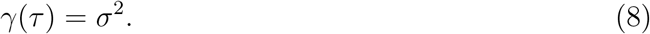

The IID model is unlikely in most data, however it does serve as a null model against which comparisons can be made. For instance, if only a small number of distantly related species are being compared, there may be no statistically detectable phylogenetic autocorrelation in the trait data.

### BM: Infinitely diffusive evolutionary process

Brownian Motion (BM) describes a random walk with a fixed mean and infinite variance. Its semi-variance function is given by

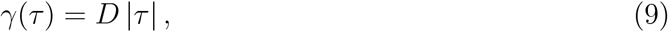

where D is the diffusion rate of the process and is proportional to the variance that can be expected over any given time lag. Under a BM model of evolution, traits are free to evolve without constraint. The semi-variance function, therefore, increases without bound. BM is likely most relevant when studying highly plastic traits, or comparing traits across taxa that have only recently diverged.

### OU: Diffusive evolutionary process with mean reversion

The OU process generalizes Brownian motion by specifying an upper limit on the process, *σ*^2^, centered around a mean. Within the bounds of *σ*^2^, the trait undergoes a random search for an optimal value but with a tendency to stay near the mean. The semi-variance function for the OU model of evolution is

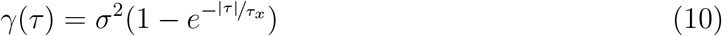

where *τ_x_* is a trait’s mean-reversion timescale, and *σ*^2^ is the variance of the process. The OU process is equivalent to the BM process in the limit of no central tendency, *τ_x_* → ∞, and infinite divergence, *σ*^2^ → ∞, with fixed diffusion rate *D* = *σ*^2^/*τ_x_*. The OU process represents a variation of Brownian diffusion when *τ* ≪ *τ_x_*, and, due to the central tendency, asymptotes to a constant variance when *τ* ≫ *τ_x_*. In other words, BM and OU can be indistinguishable over short timescales because a trait has not yet diverged sufficiently far from the mean and it is only over longer time lags that it is possible to tell the difference between BM and OU processes. In the limit where *τ* → 0, *D* = *σ*^2^/*τ_x_* → ∞ and the OU process converges to an IID model of evolution.

### Empirical case studies

#### Musteloid Sexual Size Dimorphism

We first demonstrate the utility of semi-variograms for modelling the evolution of SSD in the carnivoran superfamily Musteloidea. Musteloids are an ecologically diverse group of carnivores that exhibit substantial variability in sexual size dimorphism (SSD), ranging from parity, to males being more than twice the size of females (Noonan *et al*., 2016). Here we use a dataset describing sexual size dimorphism in 48 species of extant musteloids gathered gathered by Noonan et al. (2015). We paired these data with a time-scaled phylogenetic tree of the musteloids compiled by (Law *et al*., 2017, Fig. 3a). For these data we estimated the empirical semi-variogram using the BM weights described above, and fit IID, BM, and OU processes to the data using the methods implemented in the `R` package `slouch` (ver 2.1.4; Kopperud *et al*., 2020).

**Figure 3:**
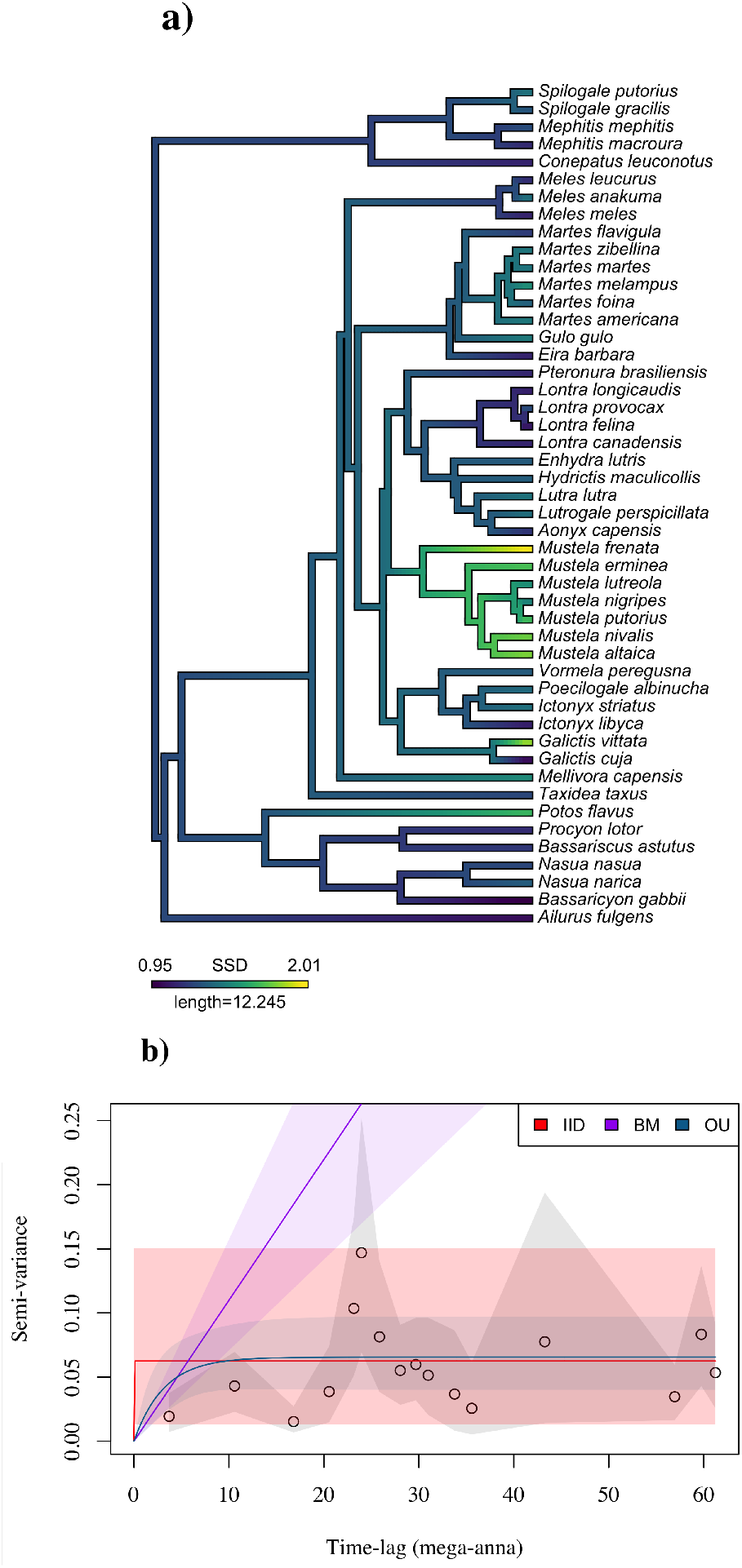
Phylogenetic tree and semi-variogram on musteloid sexual size dimorphism (SSD).

#### Carnivora maximum *per capita* rate of population growth (*r*)

We next demonstrate the utility of phylogenetic semi-variograms for modelling maximum per capita rate of population growth, *r*, in the Carnivora, a central measure of population biology. We use dataset describing *r* in 63 species of extant carnivores gathered gathered by Fagan et al. (2013). We paired these data with a phylogenetic tree of the Carnivora compiled by (Agnarsson *et al*., 2010, Fig. 4a). The *r* values were log-scaled prior to analysis and the empirical semi-variogram and model fitting process were carried out as described above.

**Figure 4:**
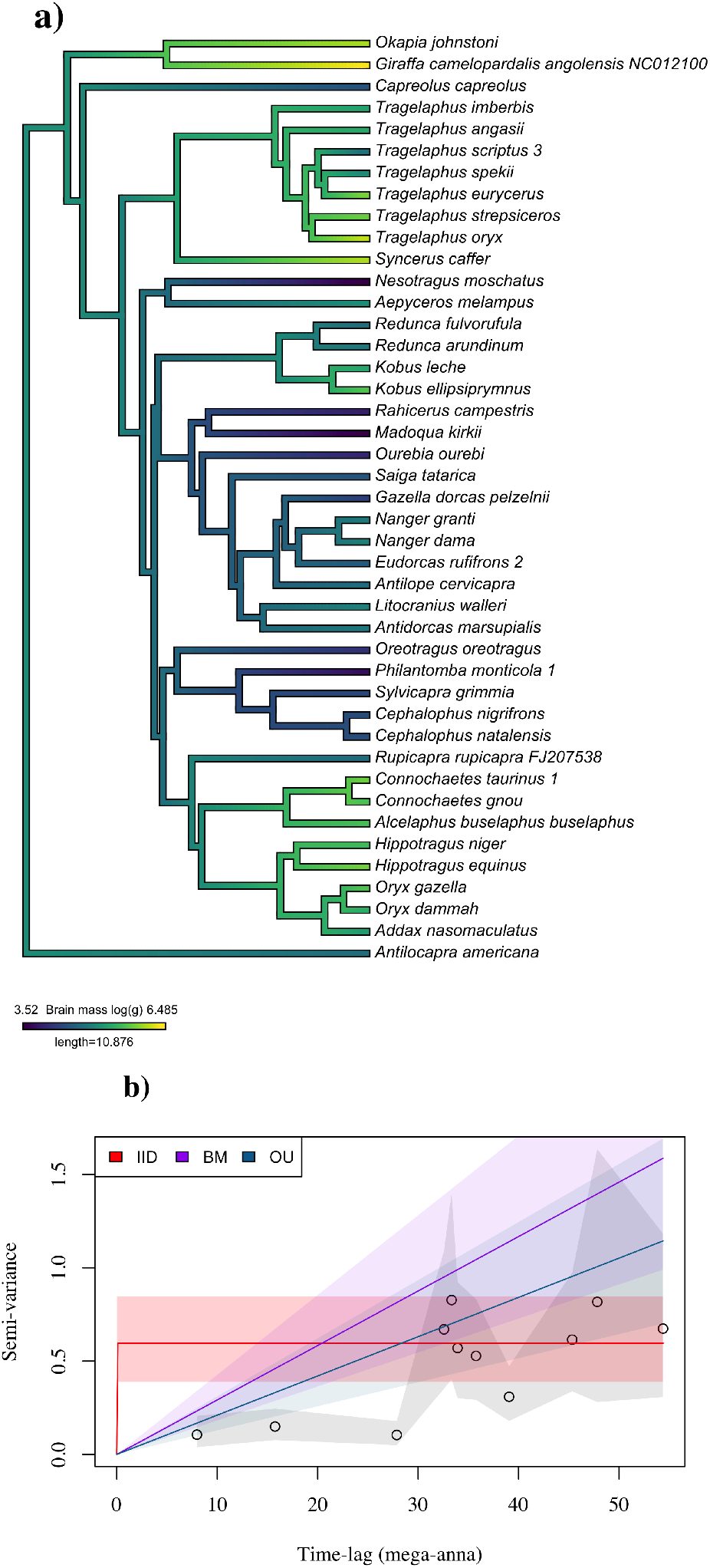
Phylogenetic tree and semi-variogram on maximum *per capita* rate of population growth, *r*, in the Carnivora.

#### Artiodactyla brain size

Finally, we apply phylogenetic semi-variograms when modelling the evolution of brain size in the Artiodactyla, which represents the even-toed ungulates. Here we used morphological data on log-scaled mean brain size (g) described in Haarmann (1975) and Oboussier (1979), and openly available in the `R` package `slouch`. We paired these data with a phylogenetic tree of the Artiodactyla compiled by (Toljagić *et al*., 2018, Fig. 5a), also available in slouch. Again, the empirical semi-variogram and model fitting process were carried out as described above.

**Figure 5:**
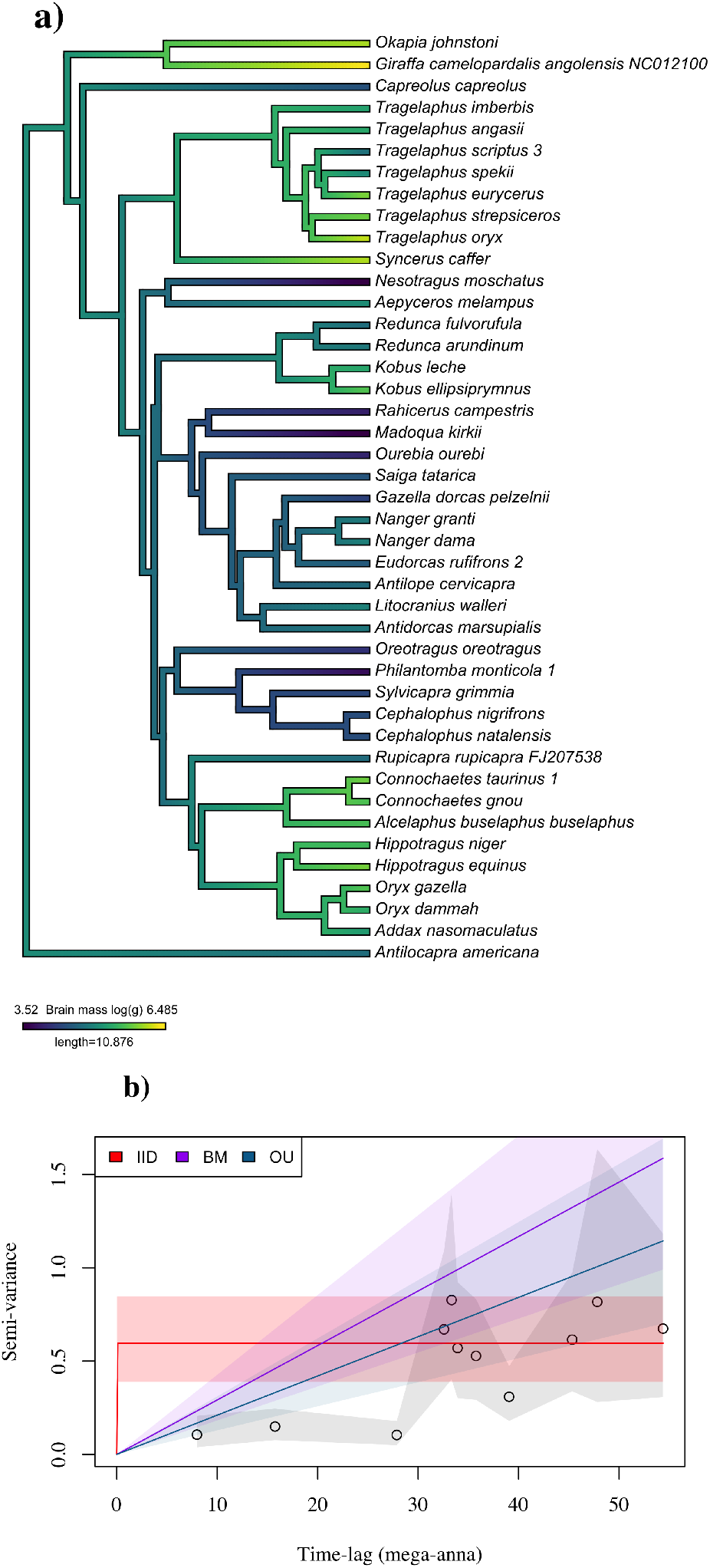
Phylogenetic trees and semi-variogram for brain size in the Artiodactyla.

The `R` script required to reproduce these analyses is presented in Appendix S1.

## Results

### Musteloid SSD

When modelling musteloid SSD, AICc based model selection identified the OU process as the best fit to the data, though with marginal support for an IID model (Table 1). There was substantially less support for the BM model. While the AICc values provided support for SSD evolving according to an OU process, they provide no information on why this model was selected over the BM or IID processes, nor on why an IID model had substantially more support than BM. Comparing the fitted models against the empirical semi-variogram demonstrates the reason for the preference of the OU model over the BM or IID processes (Fig. 3b). The empirical semi-variogram for the variance in SSD shows clear asymptotic behaviour. The infinitely diffusive BM model was the least supported based on AICc values, and the semi-variogram clearly shows this mismatch. The IID model, in contrast, captures the asymptotic behaviour of the empirical variogram, but misses the phylogenetic autocorrelation over shorter time-scales. Only the OU model captures both features of the semi-variogram.

**Table 1:**
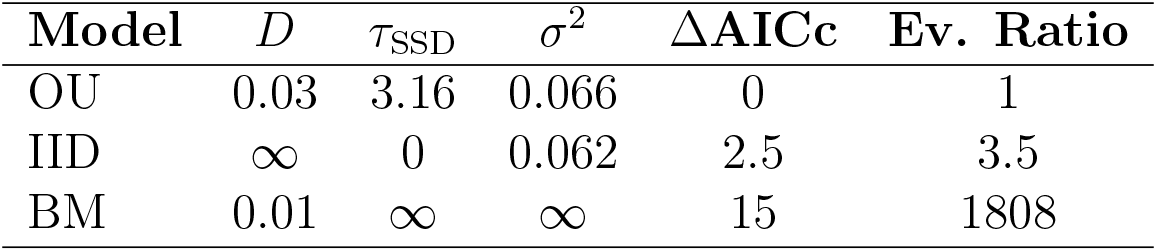
Small-sample-size corrected Akaike Information Criterion (AICc) differences for models fit to musteloid sexual size dimorphism. In the first three columns the diffusion rate (*D*, in mega-anna^-1^), phylogenetic autocorrelation timescale (*τ*_SSD_, in mega-anna), and stationary variance (*σ*^2^), are shown. The evidence ratios for each model was calculated as 1/e^-1/2ΔAICc^.

### Carnivora *r*

We found that the evolution of the intrinsic rate of increase, *r*, in the Carnivora was best described by an OU process (Table 2), with the empirical semi-variogram showing clear asymptotic behaviour (Fig. 4b). In other words, the best fit model suggests that population growth rates are not expected to evolve without bound, but instead fluctuate around a mean. Here, although the IID model was able to capture the asymptotic behaviour of the variogram, it missed the phylogenetic inertia over shorter evolutionary timescales. As a result it received ~ 1.9 × 10^12^ times less support than the OU model and ~ 7.5 × 10^11^ less support than BM.

**Table 2:** Small-sample-size corrected Akaike Information Criterion (AICc) differences for models fit to the log-scaled maximum per capita rate of population growth, *r*, in the Carnivora. In the first three columns the diffusion rate (D, in proportional units of time^-1^), phylogenetic autocorrelation timescale (*τ_r_*, in proportional units of time), and stationary variance (*σ*^2^), are shown. The evidence ratios for each model was calculated as 1/e-^1/2ΔAICc^.

| Model | $D$ | $\tau_r$ | $\sigma^2$ | $\Delta\text{AICc}$ | Ev. Ratio |
| --- | --- | --- | --- | --- | --- |
| OU | 2.84 | 0.43 | 0.87 | 0 | 1 |
| BM | 2.14 | ∞ | ∞ | 2 | 2.7 |
| IID | ∞ | 0 | 0.94 | 56.6 | $1.9 \times 10^{12}$ |

### Artiodactyla brain size

For brain size in the Artiodactyla, AICc based model selection identified the BM process as the best fit to the data, though with marginal support for an OU model, and substantially less support for the IID model (Table 3). Again, while the AICc values provided support for brain size evolving according to a BM process, they provide no information on why this model was selected over the OU or IID processes. Comparing the fitted models against the empirical semi-variogram demonstrates the reason for the preference of the BM model over the OU or IID processes (Fig. 5b). In contrast to the previous examples, here the empirical semi-variogram shows no clear asymptotic behaviour. In other words, variance in brain size was simply proportional to phylogenetic distance. The infinitely diffusive BM model was therefore the most appropriate model for the data, and the semi-variogram shows this correspondence. Thus, even though the OU variogram falls visually closer to the empirical data (Fig. 5b), it is not supported because the fit requires an additional parameter.

**Table 3:** Small-sample-size corrected Akaike Information Criterion (AICc) differences for models fit to artiodactyla brain size data. In the first three columns the diffusion rate (D, in mega-anna^-1^), phylogenetic autocorrelation timescale (*τ_r_*, in mega-anna), and stationary variance (*σ*^2^), are shown. The evidence ratios for each model was calculated as 1/e-^1/2ΔAICc^.

| Model | $D$ | $\tau_r$ | $\sigma^2$ | $\Delta\text{AICc}$ | Ev. Ratio |
| --- | --- | --- | --- | --- | --- |
| BM | 0.03 | ∞ | ∞ | 0 | 0 |
| OU | 0.03 | $4.3 \times 10^5$ | $9.1 \times 10^3$ | 2.4 | 3.3 |
| IID | ∞ | 0 | 0.77 | 31.3 | $6.3 \times 10^6$ |

## Discussion

Since Felsenstein (1985) translated the concept of phylogenetic inertia into a statistical problem, methods for modelling evolutionary processes have been advancing rapidly (e.g., Martins & Hansen, 1997; Abouheif, 1999; Butler & King, 2004; Harmon *et al*., 2008; Revell *et al*., 2008; Blomberg *et al*., 2020), allowing for substantially more profound insight into evolutionary biology than was previously possible. While these modelling approaches all treat phylogenetic inertia as a form of statistical autocorrelation, no tools exist to visualise the autocorrelation structure of the data. As such, researchers often rely on colouring the branch tips or lengths of phylogenetic trees based on trait values (e.g., Revell, 2013). While this can serve as a useful visual tool, it provides no information on the underlying stochastic process by which the trait may be evolving, nor on whether any of the candidate models actually look like the data. Here we extended semi-variograms developed for visualizing temporal autocorrelation in other contexts (Fleming *et al*., 2014) to the needs of phylogenetic autocorrelation, and show how phylogenetic variograms can be used to straightforwardly visualise phylogenetic autocorrelation structures. Notably, because the variograms of the IID, BM, and OU processes each have different theoretical profiles, comparing fitted semi-variance functions against empirical semi-variograms can serve as a useful diagnostic tool, allowing researchers to understand why any given evolutionary model might be selected over another, which features are well captures by the model, and which are not.

When modelling musteloid SSD, the empirical variogram showed a tendency for the variance in SSD to stabilise over time, a characteristic feature of OU evolution. In agreement with this visual assessment, AIC-based model selection identified the OU model as the best fit to the data. In contrast, the BM model received substantially less support than the OU model. While AIC-based model selection yielded a clear ‘top’ model, without the empirical variogram it would not have been possible to see that the failure of the infinitely diffusive BM model was because it did not capture the asymptotic behaviour of the data. Indeed, for musteloid SSD a white noise, IID process was actually a better fit to the variogram than the BM model. The phylogenetic variogram proved equally useful in understanding why the OU model was selected when modelling population growth rate in the Carnivora, and why BM was the selected evolutionary model for brain size in the Artiodactyla. For the time-scale over which species in the Artiodactyla have been diverging, the variance in brain size has yet to show any evidence of stabilising selection (Fig. 5b). Here BM showed good correspondence with the empirical variogram.

It is important to note that while variograms can serve as valuable diagnostic tools, they are only as useful as the pool of models against which they can be compared. In other words, it is entirely possible that none of the candidate models end up being a good match to the data. While variograms on their own do nothing to solve this problem, they do allow research to identify potential pitfalls and areas for model improvement. For instance, the empirical variograms for both growth rate in the Carnivora, and brain size in the Artiodactyla showed strong phylogenetic inertia over shorter timescales that was not well captured by any of the models fitted here (Figs. 4b, and 5b). A model with multiple autocorrelation timescales would likely be a better match to those data (e.g., Johnson *et al*., 2008; Fleming *et al*., 2014), and is a promising area of future research. In this regard, however, it is also important to note that variogram errors are correlated (Diggle *et al*., 1998), and smooth trends may not necessarily be significant and should be treated with caution. Finally, another important limitation of phylogenetic variograms is that the time-lags for phylogenetic variograms are calculated based on the topologies and branch lengths of the supplied phylogenetic trees. As such, the structure of a particular tree will dictate what pairwise time-lags are possible and the resulting time series will almost always be heavily irregular. When combined with pronounced interspecific variability and small sample sizes, phylogenetic variograms can be difficult to interpret. In addition, although we did not assess the impact of tree topology on parameter estimation here, the model fitting process can be strongly affected by the data density across the time lags. Despite these limitations, however, phylogenetic variograms can serve as a useful tool for visualising the autocorrelation structure of evolutionary processes, and informing future model developments.

When working with autocorrelated data, it is generally recommended that any analysis begin with a non-parametric estimate of the autocorrelation structure of the data that can be visualised. Although this is a key step in the data analysis ‘pipeline’, it is one that has heretofore not been possible for phylogenetic data (but see Diniz-Filho, 2001). The variograms methods developed here enable phylogenetic autocorrelation to be visualised, allowing for fitted models to be compared against the empirical variogram, facilitating model identification prior to subsequent analyses. We therefore recommend that any phylogenetic analysis begin with variogram estimation and visualisation (see also Pérez-Barbería *et al*., 2004). The methods developed in this work are openly accessible in the `R` package `ctpm`, available at https://github.com/NoonanM/ctpm.

## Author contributions

MJN, WFF, and CHF contributed to the original project conception and development. MJN developed the `ctpm R` package, analysed the data, and drafted the manuscript. CHF derived the weights and aided with the development of the ctpm package. All authors contributed to the writing.

## Data Availability

All data used in the manuscript are currently openly accessible. The musteloid trait data were obtained from Noonan et al. (2015) and the phylogenetic tree from Law et al. (2017). The data on maximum *per capita* rate of population growth, *r*, in the Carnivora were obtained from Fagan et al. (2013) and the phylogenetic tree of the Carnivora from Agnarsson et al. (2010). The data on log-scaled mean brain size and the phylogenetic tree of the Artiodactyla are openly available in the `R` package `slouch` (Toljagić *et al*., 2018).

## Notes

### Competing Interest Statement

The authors have declared no competing interest.

## References

Abouheif, E. & Fairbairn, D.J. (1997) A comparative analysis of allometry for sexual size dimorphism: assessing Rensch’s rule. American Naturalist.

Abouheif, E. (1999) A method for testing the assumption of phylogenetic independence in comparative data. Evolutionary Ecology Research, 1, 895–909.

Agnarsson, I., Kuntner, M. & May-Collado, L.J. (2010) Dogs, cats, and kin: A molecular species-level phylogeny of Carnivora. Molecular Phylogenetics and Evolution, 54, 726–745.

Bergman, C. (1848) Uber die Verhaltnisse der Warmeokonomie der Thiere zu ihrer Grosse. Göttinger Studien, Göttingen.

Blomberg, S.P. & Garland, T. Jr (2002) Tempo and mode in evolution: phylogenetic inertia, adaptation and comparative methods. Journal of Evolutionary Biology, 15, 899–910.

Blomberg, S.P., Garland, T.Jr & Ives, A.R. (2003) Testing for phylogenetic signal in comparative data: behavioral traits are more labile. Evolution, 57, 717–745.

Blomberg, S.P., Rathnayake, S.I. & Moreau, C.M. (2020) Beyond Brownian motion and the Ornstein-Uhlenbeck process: Stochastic diffusion models for the evolution of quantitative characters. The American Naturalist, 195, 145–165.

Brown, J.H., Gillooly, J.F., Allen, A.P., Savage, V.M. & West, G.B. (2004) Toward a metabolic theory of ecology. Ecology, 85, 1771–1789.

Burnham, K.P. & Anderson, D.R. (2002) Model Selection and Multimodel Inference: A Practical Information-Theoretic Approach. Springer New York, New York, NY.

Butler, M.A. & King, A.A. (2004) Phylogenetic comparative analysis: a modeling approach for adaptive evolution. The American Naturalist, 164, 683–695.

Clavel, J., Escarguel, G. & Merceron, G. (2015) mvMORPH: an R package for fitting multivariate evolutionary models to morphometric data. Methods in Ecology and Evolution, 6, 1311–1319.

Darwin, C. (1859) On the origin of species by means of natural selection. John Murray, London.

Diggle, P.J., Tawn, J.A. & Moyeed, R.A. (1998) Model-based geostatistics. Journal of the Royal Statistical Society: Series C (Applied Statistics), 47, 299–350.

Diniz-Filho, J.A.F. (2001) Phylogenetic autocorrelation under distinct evolutionary processes. Evolution, 55, 1104–1109.

Einstein, A. (1905) Zur Elektrodynamik bewegter Körper. (German) [On the electrodynamics of moving bodies]. Annalen der Physik, 322, 891–921.

Fagan, W.F., Pearson, Y.E., Larsen, E.A., Lynch, H.J., Turner, J.B., Staver, H., Noble, A.E., Bewick, S. & Goldberg, E.E. (2013) Phylogenetic prediction of the maximum per capita rate of population growth. Proceedings of the Royal Society B: Biological Sciences, 280, 20130523.

Felsenstein, J. (1985) Phylogenies and the comparative method. The American Naturalist, 125, 1–15.

Fleming, C.H., Calabrese, J.M., Mueller, T., Olson, K.A., Leimgruber, P. & Fagan, W.F. (2014) From Fine-Scale Foraging to Home Ranges: A Semivariance Approach to Identifying Movement Modes across Spatiotemporal Scales. The American Naturalist, 183, E154–E167.

Furness, A.I., Avise, J.C., Pollux, B.J., Reynoso, Y. & Reznick, D.N. (2021) The evolution of the placenta in poeciliid fishes. Current Biology, 31, 2004–2011.e5.

Garland, T., Bennett, A.F. & Rezende, E.L. (2005) Phylogenetic approaches in comparative physiology. Journal of experimental Biology, 208, 3015–3035.

Haarmann, K. (1975) Morphological and histological study of neocortex of bovides (antilopinae, cephalophinae) and tragulidae with comments on evolutionary development. Journal Fur Hirnforschung, 16, 93–116.

Harmon, L.J., Weir, J.T., Brock, C.D., Glor, R.E. & Challenger, W. (2008) Geiger: investigating evolutionary radiations. Bioinformatics, 24, 129–131.

Harvey, P.H. & Pagel, M.D. (1991) The comparative method in evolutionary biology. Oxford.

Herrera, C.M. (2020) Flower traits, habitat, and phylogeny as predictors of pollinator service: a plant community perspective. Ecological Monographs, 90, e01402.

Hirt, M.R., Jetz, W., Rall, B.C. & Brose, U. (2017) A general scaling law reveals why the largest animals are not the fastest. Nature Ecology & Evolution, 1, 1116.

Jetz, W. (2004) The scaling of animal space use. Science, 306, 266–268.

Johnson, D.S., London, J.M., Lea, M.A. & Durban, J.W. (2008) Continuous-time correlated random walk model for animal telemetry data. Ecology, 89, 1208–1215.

Johnson, P.J., Noonan, M.J., Kitchener, A.C., Harrington, L.A., Newman, C. & Macdonald, D.W. (2017) Rensching cats and dogs: Feeding ecology and fecundity trends explain variation in the allometry of sexual size dimorphism. Royal Society Open Science, 4.

Kellermann, V., Loeschcke, V., Hoffmann, A.A., Kristensen, T.N., Fløjgaard, C., David, J.R., Svenning, J.C. & Overgaard, J. (2012) Phylogenetic constraints in key functional traits behind species’climate niches: Patterns of desiccation and cold resistance across 95 Drosophila species. Evolution: International Journal of Organic Evolution, 66, 3377–3389.

Kopperud, B., Pienaar, J., Voje, K., Orzack, S. & Hansen, T. (2020) slouch: Stochastic Linear Ornstein-Uhlenbeck Comparative Hypotheses. R package version.

Law, C.J., Slater, G.J. & Mehta, R.S. (2017) Lineage diversity and size disparity in Musteloidea: Testing patterns of adaptive radiation using molecular and fossil-based methods. Systematic Biology, 67, 127–144.

Lukas, D. & Clutton-Brock, T.H. (2013) The evolution of social monogamy in mammals. Science, 341, 526–530.

MacQueen, J. et al. (1967) Some methods for classification and analysis of multivariate observations. Proceedings of the fifth Berkeley symposium on mathematical statistics and probability, volume 1, pp. 281–297. Oakland, CA, USA.

Martins, E.P. & Hansen, T.F. (1997) Phylogenies and the comparative method: a general approach to incorporating phylogenetic information into the analysis of interspecific data. The American Naturalist, 149, 646–667.

Matheron, G. (1963) Principles of geostatistics. Economic geology, 58, 1246–1266.

McLachlan, G.J. & Basford, K.E. (1988) Mixture models: Inference and applications to clustering, volume 38. M. Dekker New York.

Morales, E. (2000) Estimating phylogenetic inertia in Tithonia (Asteraceae): a comparative approach. Evolution, 54, 475–484.

Moran, P.A. (1950) Notes on continuous stochastic phenomena. Biomeetrika, 37, 17–23.

Mouselimis, L. (2021) ClusterR: Gaussian Mixture Models, K-Means, Mini-Batch-Kmeans, K-Medoids and Affinity Propagation Clustering.

Münkemüller, T., Lavergne, S., Bzeznik, B., Dray, S., Jombart, T., Schiffers, K. & Thuiller, W. (2012) How to measure and test phylogenetic signal. Methods in Ecology and Evolution, 3, 743–756.

Noonan, M.J., Fleming, C.H., Tucker, M.A., Kays, R., Harrison, A.L., Crofoot, M.C., Abrahms, B., Alberts, S.C., Ali, A.H., Altmann, J. et al. (2020) Effects of body size on estimation of mammalian area requirements. Conservation Biology, 34, 1017–1028.

Noonan, M.J., Johnson, P.J., Kitchener, A.C., Harrington, L.A., Newman, C. & Macdonald, D.W. (2016) Sexual size dimorphism in musteloids: An anomalous allometric pattern is explained by feeding ecology. Ecology and Evolution, 6, 8495–8501.

Noonan, M.J., Newman, C., Buesching, C.D. & Macdonald, D.W. (2015) Evolution and function of fossoriality in the Carnivora: Implications for group-living. Frontiers in Ecology and Evolution, 3, 726.

Oboussier, H. (1979) Evolution of the brain and phylogenetic development of Mrican Bovidae. African Zoology, 14, 119–124.

Pagel, M. (1999) Inferring the historical patterns of biological evolution. Nature, 401, 877–884.

Pérez-Barbería, F., Elston, D., Gordon, I. & Illius, A. (2004) The evolution of phylogenetic differences in the efficiency of digestion in ruminants. Proceedings of the Royal Society of London Series B: Biological Sciences, 271, 1081–1090.

Rensch, B. (1950) Die abhängigkeit der relativen sexualdifferenz von der körpergrösse. Bonner Zoologische Beiträge, 1, 58–69.

Revell, L.J. (2012) phytools: an R package for phylogenetic comparative biology (and other things). Methods in Eecology and Evolution, 3, 217–223.

Revell, L.J. (2013) Two new graphical methods for mapping trait evolution on phylogenies. Methods in Ecology and Evolution, 4, 754–759.

Revell, L.J., Harmon, L.J. & Collar, D.C. (2008) Phylogenetic signal, evolutionary process, and rate. Systematic Biology, 57, 591–601.

Smaers, J.B., Rothman, R.S., Hudson, D.R., Balanoff, A.M., Beatty, B., Dechmann, D.K.N., de Vries, D., Dunn, J.C., Fleagle, J.G., Gilbert, C.C., Goswami, A., Iwaniuk, A.N., Jungers, W.L., Kerney, M., Ksepka, D.T., Manger, P.R., Mongle, C.S., Rohlf, F.J., Smith, N.A., Soligo, C., Weisbecker, V. & Safi, K. (2021) The evolution of mammalian brain size. Science Advances, 7.

Toljagić, O., Voje, K.L., Matschiner, M., Liow, L.H. & Hansen, T.F. (2018) Millions of years behind: slow adaptation of ruminants to grasslands. Systematic Biology, 67, 145–157.

Turlach, M.B.A. (2019) Package ‘quadprog’.

Uhlenbeck, G.E. & Ornstein, L.S. (1930) On the theory of the Brownian motion. Physical review, 36, 823–841.

