## Appendix S1 for "A semi-variance approach to visualising phylogenetic autocorrelation": Appendix_S1.html

Appendix S1 - Empirical examples


### Appendix S1 - Empirical examples

###### Noonan MJ, Fagan WF, Fleming CH

This file was last updated on May 21, 2021.

---

#### Overview

In this appendix we detail the application of phylogenetic semi-variograms to three empirical datasets to arrive at the results presented in the main text.

---

#### Musteloid Sexual Size Dimorphism

We first demonstrate the utility of semi-variograms for modelling the evolution of SSD in the carnivoran superfamily Musteloidea. Musteloids are an ecologically diverse group of carnivores that exhibit substantial variability in SSD, ranging from parity, to males being more than twice the size of females (Noonan et al. 2016). Here we use a dataset describing sexual size dimorphism in 48 species of extant musteloids gathered gathered by (Noonan et al. 2015). We paired these data with a time-scaled phylogenetic tree of the musteloids compiled by (Law, Slater, and Mehta 2017). For these data we estimated the empirical semi-variogram using the BM weights described above, and fit IID, BM, and OU processes to the data using the methods implemented in the `R` package `slouch` (Kopperud et al. 2020).

```
# Load in the requisite packages
library(viridis)
library(phytools)
library(ctpm)
library(slouch)

#####
#Load in the data and the tree
data("moid_traits")
data("musteloids")

SSD <- moid_traits$SSD
names(SSD) <- moid_traits$Binomial

#Calculate the variogram
SVF <- variogram(SSD, musteloids, progress = F)

#Fit the evolutionary models
IID.FIT <- ctpm.fit(SSD, musteloids, model = "IID")
BM.FIT <- ctpm.fit(SSD, musteloids, model = "BM")
OU.FIT <- ctpm.fit(SSD, musteloids, model = "OU")


#Plot the phylogenetic tree
#Use contMap from phytools
COLS2 <- viridis(length(SSD))
tree.trait <- contMap(musteloids, SSD, plot = F)
tree.trait<-setMap(tree.trait, colors=viridis(length(SSD)))

plot.contMap(tree.trait,
             type = "phylogram",
             legend = 0.4*max(nodeHeights(musteloids)),
             fsize = c(0.5, 0.6),
             res = 1000,
             lwd=2,
             outline = T,
             leg.txt = "SSD")
```

```
##############
# Now plot the variograms and fitted models
plot(SVF,
     list(IID.FIT,
          BM.FIT,
          OU.FIT),
     col.CTPM = c("red",
                  "purple",
                  "#046C9A"))
legend("topleft",
       fill = c("red",
                "purple",
                "#046C9A"),
       legend = c("IID",
                  "BM",
                  "OU"),
       horiz = T,
       cex = 0.8)
```

---

#### Carnivora maximum *per capita* rate of population growth

We next demonstrate the utility of phylogenetic semi-variograms for modelling maximum per capita rate of population growth, \(r\), in the Carnivora, a central measure of population biology. We use dataset describing \(r\) in 63 species of extant carnivores gathered gathered by (Fagan et al. 2013). We paired these data with a phylogenetic tree of the Carnivora compiled by (Agnarsson, Kuntner, and May-Collado 2010). The \(r\) values were log-scaled prior to analysis and the empirical semi-variogram and model fitting process were carried out as described above.

```
DATA <- read.csv("Data/Growth_Rate.csv")
tree <- ape::read.nexus("Data/Fagan_etal.txt")


'%ni%' <- Negate('%in%')
tree <- drop.tip(tree, c(which(tree$tip.label %ni% DATA$Species)))
row.names(DATA) <-  DATA$Species
DATA <- DATA[match(tree$tip.label, DATA[,"Species"]),]


#Data preparation
r <- log(DATA$r)
names(r) <- DATA$Species

#Calculate the variogram for population growth rate
SVF <- variogram(r, tree, trait.units = "f", progress = F, time.units = "unknown")

#Fit the evolutionary models to 
IID.FIT <- ctpm.fit(r, tree, model = "IID", time.units = "unknown")
BM.FIT <- ctpm.fit(r, tree, model = "BM", time.units = "unknown")
OU.FIT <- ctpm.fit(r, tree, model = "OU", time.units = "unknown")


#Plot the phylogenetic tree
COLS2 <- viridis(length(r))
tree.trait <- contMap(tree, r, plot = F)
tree.trait<-setMap(tree.trait, colors=viridis(length(r)))

plot.contMap(tree.trait,
             type = "phylogram",
             legend = 0.4*max(nodeHeights(tree)),
             fsize = c(0.5, 0.6),
             res = 1000,
             lwd=2,
             outline = T,
             leg.txt = "log(r)")
```

```
##############
# Now plot the variograms and fitted models
plot(SVF,
     list(IID.FIT,
          BM.FIT,
          OU.FIT),
     col.CTPM = c("red",
                  "purple",
                  "#046C9A"))
legend("topleft",
       fill = c("red",
                "purple",
                "#046C9A"),
       legend = c("IID",
                  "BM",
                  "OU"),
       horiz = T,
       cex = 0.8)
```

---

#### Artiodactyla brain size

Finally, we apply phylogenetic semi-variograms when modelling the evolution of brain size in the Artiodactyla, which represents the even-toed ungulates. Here we used morphological data on log-scaled mean brain size (g) described in (Haarmann 1975) and (Oboussier 1979), and openly available in the `R` package `slouch`. We paired these data with a phylogenetic tree of the Artiodactyla compiled by (Toljagić et al. 2018), also available in `slouch`. Again, the empirical semi-variogram and model fitting process were carried out as described above.

```
#####
#Load in the data and the tree
data(artiodactyla)
data(neocortex)

neocortex <- neocortex[match(artiodactyla$tip.label, neocortex[,"species"]),]
'%ni%' <- Negate('%in%')
artiodactyla <- drop.tip(artiodactyla, c(which(artiodactyla$tip.label %ni% neocortex$species)))
row.names(neocortex) <-  neocortex$species

#Data preparation
BRAIN <- neocortex$brain_mass_g_log_mean
names(BRAIN) <- neocortex$species

#Calculate the variogram for brain size
SVF <- variogram(BRAIN, artiodactyla, trait.units = "m", progress = F)

#Fit the evolutionary models for brain size
IID.FIT <- ctpm.fit(BRAIN, artiodactyla, model = "IID")
BM.FIT <- ctpm.fit(BRAIN, artiodactyla, model = "BM")
OU.FIT <- ctpm.fit(BRAIN, artiodactyla, model = "OU")


#Plot the phylogenetic tree
COLS2 <- viridis(length(BRAIN))
tree.trait <- contMap(artiodactyla, BRAIN, plot = F)
tree.trait<-setMap(tree.trait, colors=viridis(length(BRAIN)))

plot.contMap(tree.trait,
             type = "phylogram",
             legend = 0.4*max(nodeHeights(artiodactyla)),
             fsize = c(0.4, 0.5),
             res = 1000,
             lwd=2,
             outline = T,
             leg.txt = "Brain mass log(g)")
```

```
##############
# Now plot the variograms and fitted models
plot(SVF,
     list(IID.FIT,
          BM.FIT,
          OU.FIT),
     col.CTPM = c("red",
                  "purple",
                  "#046C9A"))
legend("topleft",
       fill = c("red",
                "purple",
                "#046C9A"),
       legend = c("IID",
                  "BM",
                  "OU"),
       horiz = T,
       cex = 0.8)
```

---

#### References

Agnarsson, Ingi, Matjaž Kuntner, and Laura J May-Collado. 2010. “Dogs, cats, and kin: A molecular species-level phylogeny of Carnivora.” *Molecular Phylogenetics and …* 54 (3): 726–45.

Fagan, William F., Yanthe E. Pearson, Elise A. Larsen, Heather J. Lynch, Jessica B. Turner, Hilary Staver, Andrew E. Noble, Sharon Bewick, and Emma E. Goldberg. 2013. “Phylogenetic Prediction of the Maximum <i>per Capita</i> Rate of Population Growth.” *Proceedings of the Royal Society B: Biological Sciences* 280 (1763): 20130523. https://doi.org/10.1098/rspb.2013.0523.

Haarmann, K. 1975. “Morphological and Histological Study of Neocortex of Bovides (Antilopinae, Cephalophinae) and Tragulidae with Comments on Evolutionary Development.” *Journal Fur Hirnforschung* 16 (2): 93–116.

Kopperud, BT, J Pienaar, KL Voje, SH Orzack, and TF Hansen. 2020. “Slouch: Stochastic Linear Ornstein-Uhlenbeck Comparative Hypotheses.” *R Package Version*, no. 2.1.4.

Law, Chris J., Graham J. Slater, and Rita S. Mehta. 2017. “Lineage Diversity and Size Disparity in Musteloidea: Testing Patterns of Adaptive Radiation Using Molecular and Fossil-Based Methods.” *Systematic Biology* 67 (1): 127–44. https://doi.org/10.1093/sysbio/syx047.

Noonan, Michael J, Paul J Johnson, Andrew C Kitchener, Lauren A Harrington, Chris Newman, and David Whyte Macdonald. 2016. “Sexual size dimorphism in musteloids: An anomalous allometric pattern is explained by feeding ecology.” *Ecology and Evolution* 6 (23): 8495–8501.

Noonan, Michael J, Chris Newman, Christina D Buesching, and David Whyte Macdonald. 2015. “Evolution and function of fossoriality in the Carnivora: Implications for group-living.” *Frontiers in Ecology and Evolution* 3: 726.

Oboussier, Henriette. 1979. “Evolution of the Brain and Phylogenetic Development of Mrican Bovidae.” *African Zoology* 14 (3): 119–24.

Toljagić, Olja, Kjetil L Voje, Michael Matschiner, Lee Hsiang Liow, and Thomas F Hansen. 2018. “Millions of Years Behind: Slow Adaptation of Ruminants to Grasslands.” *Systematic Biology* 67 (1): 145–57.
